# Biophysical basis for brain folding and misfolding patterns in ferrets and humans

**DOI:** 10.1101/2025.03.05.641682

**Authors:** Gary P. T. Choi, Chunzi Liu, Sifan Yin, Gabrielle Séjourné, Richard S. Smith, Christopher A. Walsh, L. Mahadevan

## Abstract

A mechanistic understanding of neurodevelopment requires us to follow the multiscale processes that connect molecular genetic processes to macroscopic cerebral cortical formations and thence to neurological function. Using magnetic resonance imaging of the brain of the ferret, a model organism for studying cortical morphogenesis, we create *in vitro* physical gel models and *in silico* numerical simulations of normal brain gyrification. Using observations of genetically manipulated animal models, we identify cerebral cortical thickness and cortical expansion rate as the primary drivers of dysmorphogenesis and demonstrate that *in silico* models allow us to examine the causes of aberrations in morphology and developmental processes at various stages of cortical ontogenesis. Finally, we explain analogous cortical malformations in human brains, with comparisons with human phenotypes induced by the same genetic defects, providing a unified perspective on brain morphogenesis that is driven proximally by genetic causes and affected mechanically via variations in the geometry of the brain and differential growth of the cortex.

**Impact statement:** Physical gel models and numerical simulations are created to study ferret brain gyrification and cortical malformations, and comparisons with human phenotypes are presented to link genetics and brain morphogenesis.

## Introduction

Understanding the growth and form of normal and abnormal cortical convolutions (gyri and sulci) is important for the study of human neurodevelopmental diseases ***Lui et al. (2011); Geschwind and Rakic (2013); Molnár et al. (2019); Del-Valle-Anton and Borrell (2022); Akula et al. (2023a)***. During early brain development, the cortical plate expands tangentially relative to the underlying white matter ***Welker (1990)***. This pattern of growth is the central cause of gyrification; indeed tangential cortical expansion creates compressive forces on the faster-growing outer layer of the cortex and tensile forces on the attached slower-growing inner layer, and the relative-growth induced forces cause cortical folding as suggested more than a century ago ***His (1868)*** and first quantitatively elucidated nearly fifty years ago ***Richman et al. (1975)***. At a molecular and cellular level, neurogenesis, neuronal migration, and neuronal differentiation all contribute to the tangential growth of the developing cortex via processes such as an increase in either the number or size of cells ***Fietz et al. (2010); Hansen et al. (2010); Borrell (2018); Van Essen (2022)***. Recent models that take these facts into account attempt to explain gyrification in terms of a simple mechanical instability, termed sulcification ***Hohlfeld and Mahadevan (2011)***, that when iterated with variations ***Tallinen et al. (2013)*** shows that tangential expansion of the gray matter constrained by the white matter can explain a range of different morphologies seen in the brains of different organisms ***Tallinen et al. (2014); Kroenke and Bayly (2018)***. Furthermore, when deployed over developmental time to simulate normal human cortical convolution, the results can capture a substantial range of features seen in normal human fetal brain morphogenesis ***Tallinen et al. (2016)***. However, these and other similar studies ***Toro and Burnod (2005); Toro et al. (2008); Herculano-Houzel (2009); Azevedo et al. (2009); Hutton et al. (2009); Nie et al. (2010); Bayly et al. (2013); Budday et al. (2014); Garcia et al. (2021); Heuer et al. (2023); Pang et al. (2023); Schwartz et al. (2023)*** do not allow us to understand malformations of cortical development (MCD), neurodevelopmental disorders that result from disrupted human cerebral cortex formation during embryonic brain development ***Desikan and Barkovich (2016)***, owing to our inability to probe the development of the human fetal brain *in utero*. In addition, MCDs are difficult to relate to the physical properties of cerebral cortical folding because of the uncertainties in defining the effects of specific human genetic abnormalities on specific cortical features such as thickness and surface area.

An alternative strategy is to turn to model organisms to study the developmental trajectory of MCDs. Since commonly used animal models such as the mouse and rat have lissencephalic cortices, the ferret, a gyrencephalic non-primate, has been favored as an experimentally tractable laboratory organism that demonstrates cortical folding patterns that are roughly similar to that observed in the human ***Neal et al. (2007); Fietz et al. (2010); Sawada and Watanabe (2012); John-*** son et al. (***2018); Gilardi and Kalebic (2021)***. Furthermore, since the process of cortical folding in the ferret is almost exclusively postnatal, with the progressive development of cortical gyri and sulci from postnatal day 0 (P0) to adolescence (Fig. 1), it is more easily observable. Finally, the ability to perform region-specific genetic manipulation of the ferret brain through *in utero* electroporation ***Masuda et al. (2015); Tabata and Nakajima (2001)*** makes the ferret an ideal system for modeling normal and abnormal neurodevelopmental processes.

**Figure 1.**
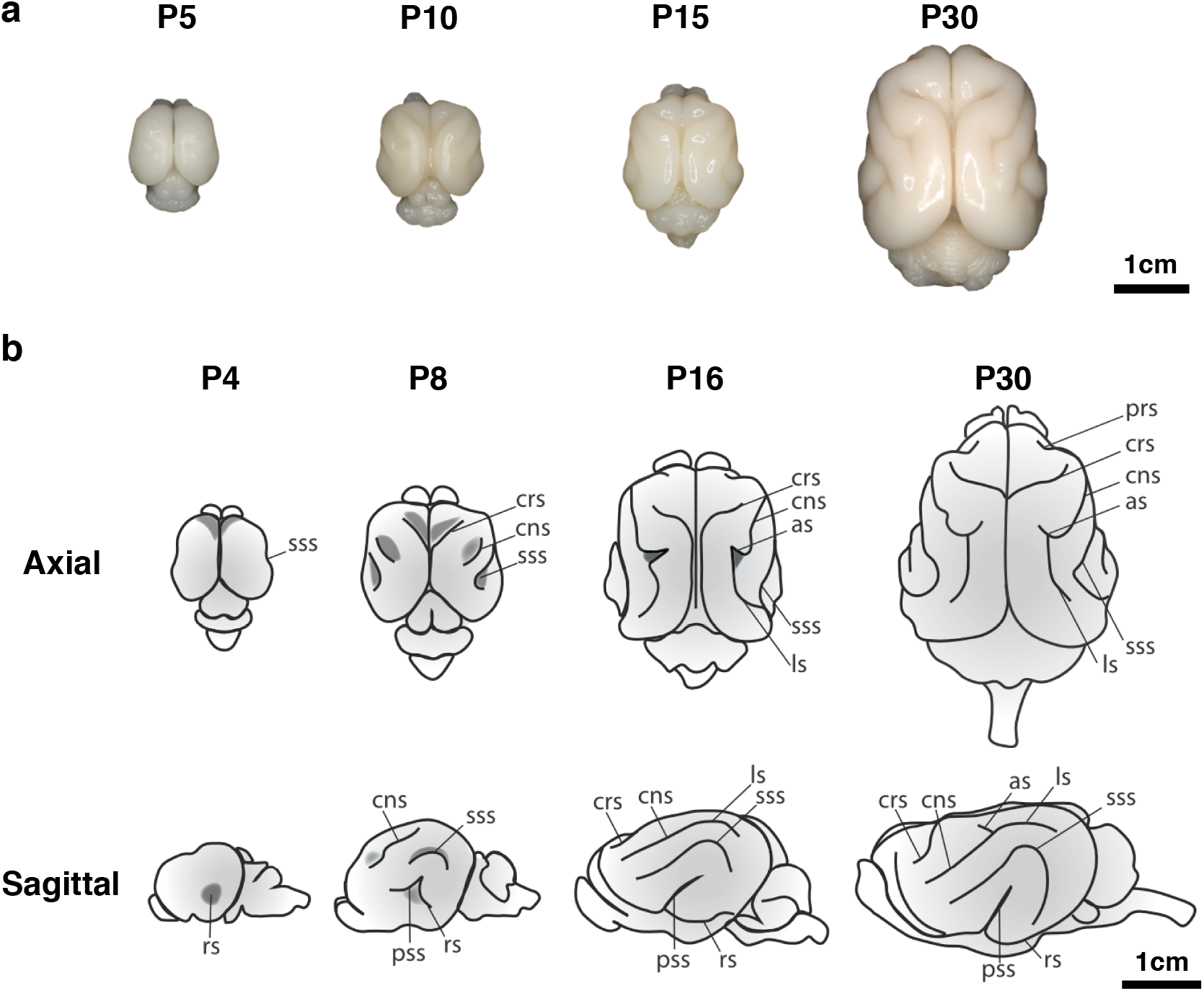
Time course of ferret brain morphogenesis. **a**, Whole brain samples from ferrets of various ages show progressive development of cortical gyri and sulci. **b**, Ferret brains show an increase in complexity of sulcal pattern and in sulcal depth throughout development. The rhinal sulcus (rs), cruciate sulcus (crs), coronal sulcus (cns), suprasylvian sulcus (sss), pseudosylvian sulcus (pss), lateral sulcus (ls), and ansate sulcus (as) are labelled. Schematic by G. Séjourné.

Inspired by our previous studies using physical experiments with swelling gels and computational models of brain growth ***Tallinen et al. (2014); Tallinen and Biggins (2015); Tallinen et al. (2016)***, we model the folding of a normal ferret brain using a physical gel model and a computational model based on the principle of constrained cortical expansion and compare the simulation results with the real brain development using various geometric morphometric approaches. We then use the computational and physical models to reproduce defective developmental processes of the ferret brain and show that they are consistent with biological experiments that manipulate different molecular drivers of neurogenesis, neuronal migration and cell growth in the cortex that underlie its relative thickness and expansion rate. Taken together, our studies provide a mesoscopic approach to brain morphogenesis that combines computational *in silico* and physical gel *in vitro* models with morphological and molecular analysis of ferret cortical disease models, and shed light on analogous MCDs in human brains.

### Physical gel model

Inspired by the observation that soft physical gels swell and fold superficially when immersed in solvents, we constructed a physical simulacrum of ferret brain folding following our previous protocols ***Tallinen et al. (2014***, 2016). Specifically, we produced two-layer PDMS gel models of the ferret brain at various ages based on surfaces reconstructed from MR Images (see SI Section S1 for details). We then immersed the two-layer gel brain model in n-hexane, which led to folding patterns by solvent-driven swelling of the outer layers (Fig. 2**a**).

**Figure 2.**
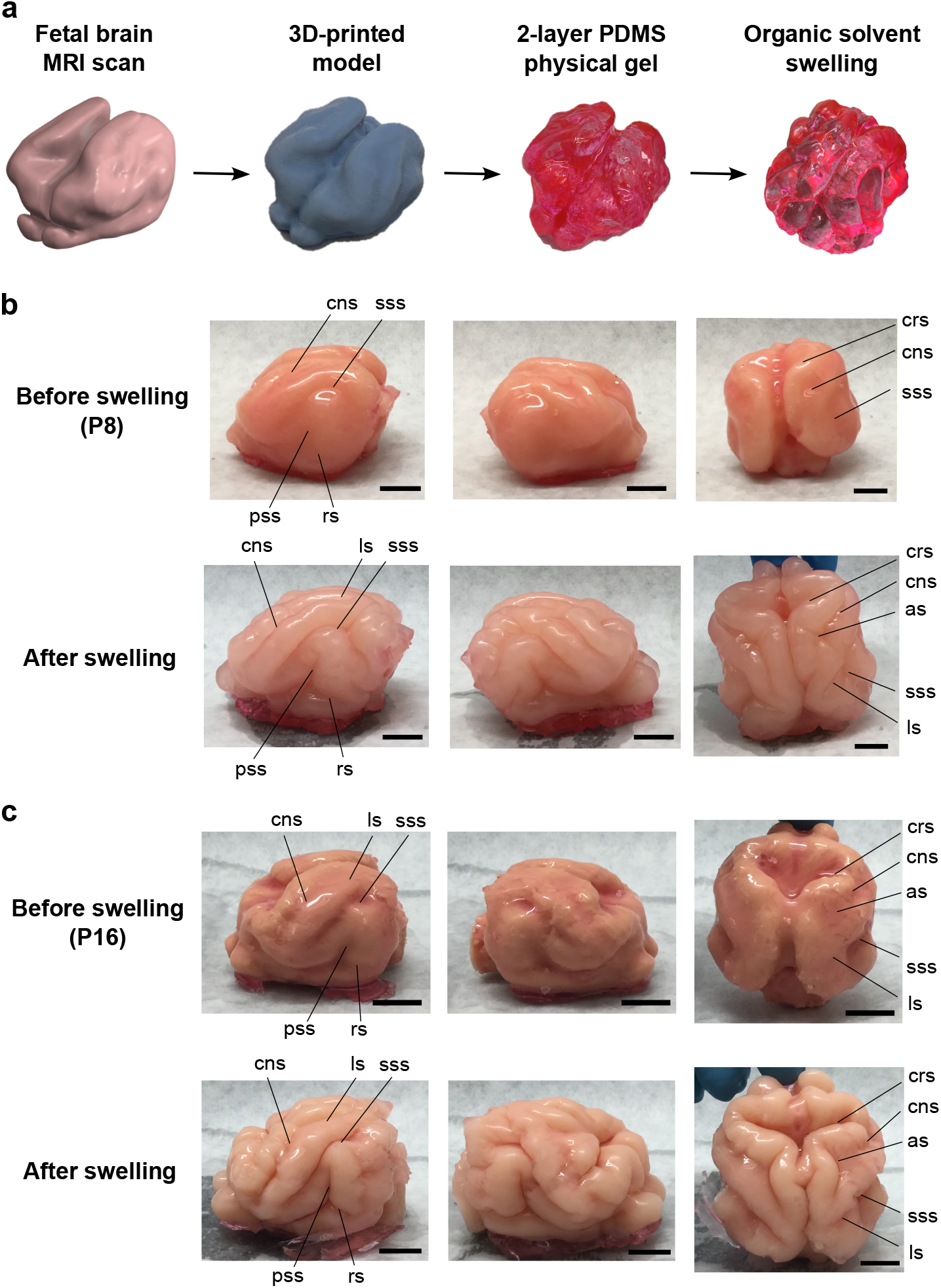
Physical gel model of ferret brain morphogenesis. **a**, Schematic of the gel experiment. We first produced a two-layer gel model of a ferret brain from MRI scans as previously described ***Tallinen et al*.** (***2016)***. We then immersed the gel model in n-hexane for 1.5 hours, which induced the outer layer to swell by absorbing the solvent over time, resulting in the development of cortical gyri and sulci. **b**, The swelling experiment for the P8 ferret model, in which it can be observed that the swelling of the cortical layer produces sulcal patterns and characteristics comparable to the real ferret brain. **c**, The swelling experiment for the P16 gel model. Scale bar = 1cm. Notation guide: cruciate sulcus (crs), coronal sulcus (cns), suprasylvian sulcus (sss), rhinal sulcus (rs), pseudosylvian sulcus (pss), lateral sulcus (ls), ansate sulcus (as).

Fig. 2**b** shows the experimental results for a P8 gel brain; it swells nonuniformly and folds progressively from an initial state that has invaginations corresponding to the cruciate sulcus (crs), the coronal sulcus (cns), and the suprasylvian sulcus (sss). The post-swelling state shows the development of sulci corresponding in location and self-contacting nature to the crs, cns, sss, and the formation of rhinal sulcus (rs), the pseudosylvian sulcus (pss), and lateral sulcus (ls), and ansate sulcus (as) observed in real ferrets aged P21 and older. Fig. 2**c** shows another swelling experiment with a starting shape corresponding to the P16 gel brain, from which we observe a similar progression in the folding patterns. We see that our minimal physical model can capture the qualitative aspects of the folding transitions in the ferret brain (see also SI Section S1 and Video S1–S2).

### Computational model

To complement our physical experiments with quantitative simulations of ferret brain development, we followed the approach in ***Tallinen et al. (2014***, 2016) and considered a neo-Hookean material model for the brain cortex consisting of a layer of gray matter on top of a deep layer of white matter with volumetric strain energy density

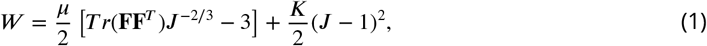

where **F** is the deformation gradient, ***J*** = det(**F**), *μ* is the shear modulus and *K* is the bulk modulus. We assume that *K* = 5*μ* for a modestly compressible material. Computer simulations were then performed on tetrahedral meshes of ferret brains to model the gyrification (see SI Section S2 for more details).

We considered both simulations that modeled the changes in brain morphology from P0 to P32 as one continuous process (Fig. 3**a**, see also SI Movie S3) and stepwise simulations that considered the growth process in stages, i.e. from P0 to P4, from P4 to P8, from P8 to P16 and from P16 to P32 (Fig. 3**b**, see also SI Movie S4). In both sets of simulations, the emergence of cortical folding can be observed. In the continuous simulation approach, we observed the appearance of multiple minor folds since the continuous simulations only depend on the P0 initial brain, so that the effect of minor features in the P0 brain on the brain growth may accumulate over time. By contrast, in the stepwise simulation approach which focuses on multiple shorter growth periods, and thus reduces the accumulation of shape variations over time. Comparing the P16 results of stepwise numerical simulation, the gel experiment and the P16 real brain, we observe that the folding patterns are visually very similar (Fig. 4**a**). For a more quantitative comparison, we applied a method based on aligning landmarks using spherical mapping termed FLASH (Fast Landmark Aligned Spherical Harmonic Parameterization) ***Choi et al. (2015)*** to parameterize the simulated P16 ferret brain and the P16 brain surface generated from the MRI scans onto the unit sphere using landmark-aligned optimized conformal mappings, with the coronal sulcus (cns), suprasylvian sulcus (sss), presylvian sulcus (prs), and pseudosylvian sulcus (pss) on both the left and right hemispheres used as landmarks (see Fig. 4**a**, and SI Section S3 for more details). We then assessed the geometric similarity of the two brain surfaces on the spherical domain in terms of their shape index ***Koenderink and van Doorn (1992)***, which is a surface measure defined based on the surface mean curvature and Gaussian curvature. The similarity of the two shape indices suggests that the folding pattern produced by our simulation is close to the actual folding pattern (see SI Section S3 for additional analyses and ***Yin et al. (2025)*** for more details of the morphometric method). We further utilized spherical harmonic-based representations for comparing the real and simulated ferret brains at different maximum orders, which also show that they have consistent geometric similarities (see SI Section S3).

**Figure 3.**
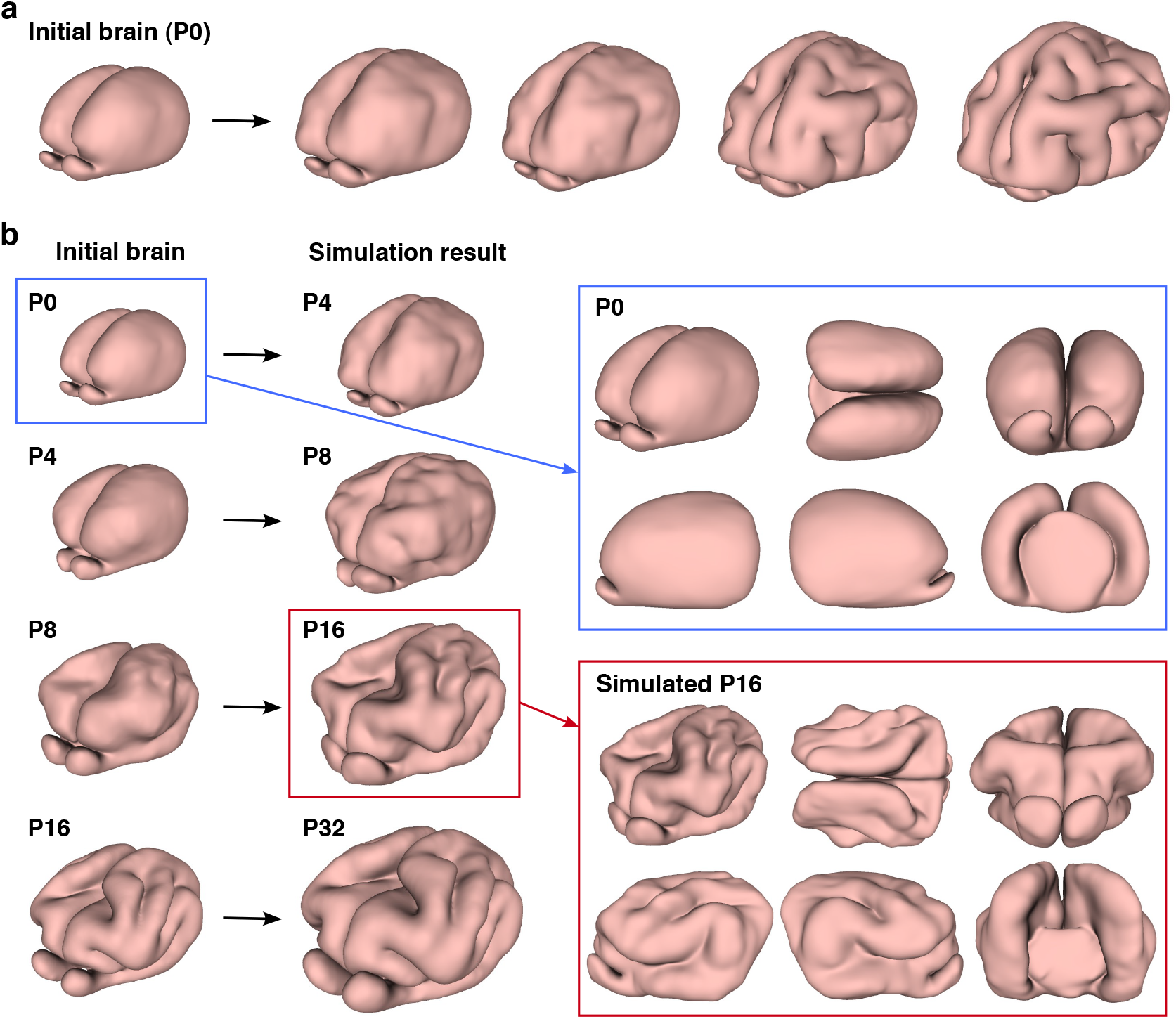
Numerical model of ferret brain morphogenesis. **a**, Continuous growth simulation from P0 to adolescence. The P0 brain tetrahedral mesh was used as the input for the numerical simulation. **b**, Stepwise growth simulation from P0 to P4, P4 to P8, P8 to P16, and P16 to P32. For different growth intervals, we use different brain tetrahedral meshes as the input for the numerical simulation. Different views of the input P0 brain and the simulated P16 brain are provided.

**Figure 4.**
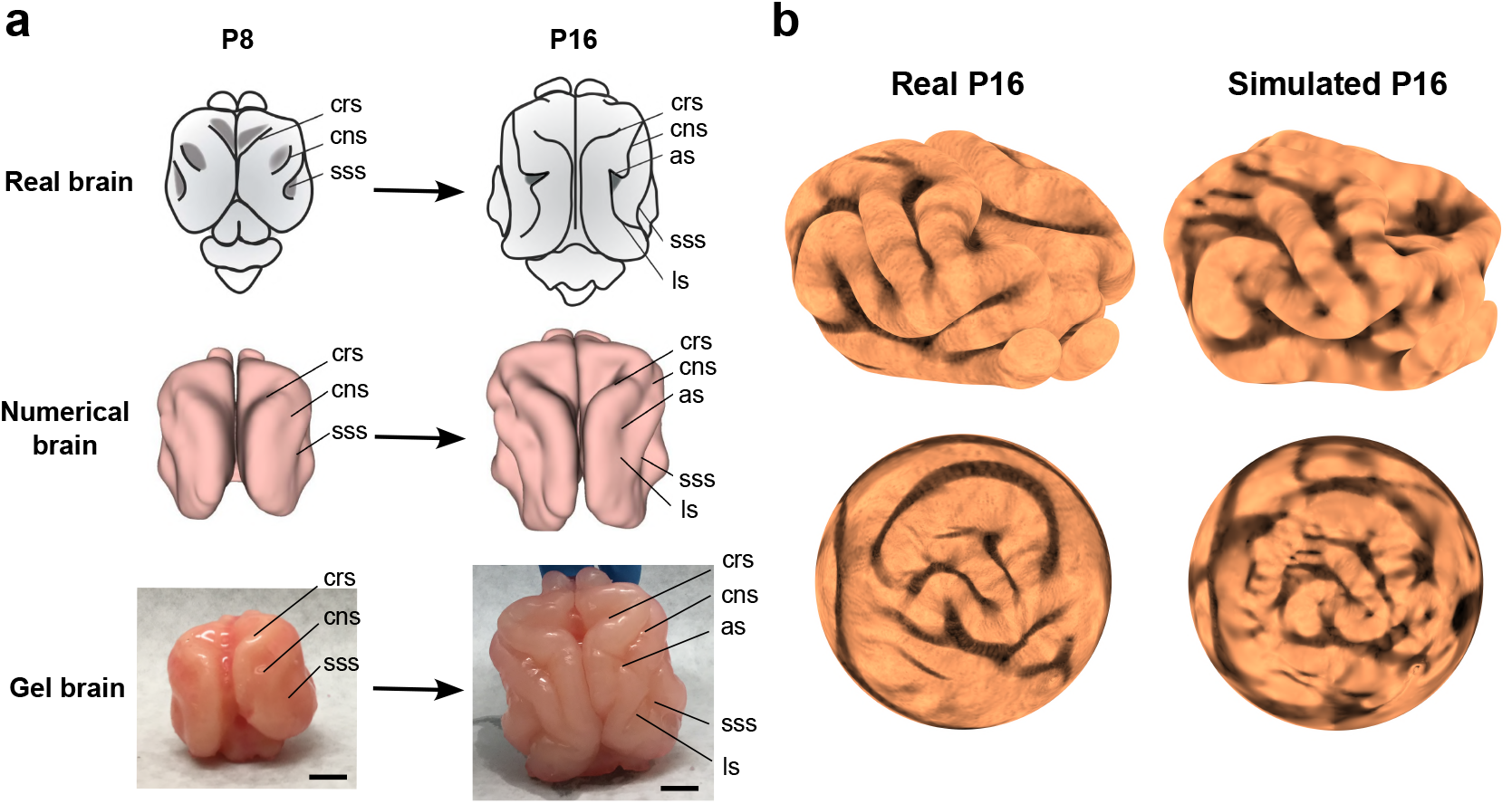
Comparison of cortical folding in real and simulated ferret brain models. **a**, The top row shows the increase in complexity of sulcal pattern and in sulcal depth of ferret brains from P8 to P16. The middle row shows a numerical model of a P8 brain and its deformed state mimicking progression to P16. The bottom row shows a physical gel model of P8 and its post-swelling state mimicking progression to P16 (scale bar = 1cm). The P8 initial states have invaginations corresponding to the cruciate sulcus (crs), coronal sulcus (cns) and suprasylvian sulcus (sss), and both the numerical deformed state and the physical post-swelling state have sulci corresponding in location and self-contacting nature to the crs, cns, sss, lateral sulcus (ls), and ansate sulcus (as) observed in P16 real ferrets. **b**, The real P16 brain reconstructed from MRI scans, the simulated P16 brain, and their respective landmark-aligned spherical mappings obtained by the FLASH algorithm ***Choi*** et al. (***2015)***, each color-coded with the shape index ***Koenderink and van Doorn (1992)*** of the brain.

### Physical gel and computational models for misfolding ferret brains

The effectiveness of our differential growth-based model in quantifying the normative development of normal ferret brains begs the question of whether we can use the same framework to study malformations of cortical development (MCD). Here, we first consider simulating ferret cortical misfolding using both modified physical gel and computational models. In Fig. 5**a**, we performed a modified numerical experiment on the P8 brain with the cortical thickness reduced to 1/4 of the original thickness globally. We also performed a modified gel brain experiment by surface-coating 1 layer of PDMS gel onto the core layer instead of 4 layers as in the original gel model in Fig. 2, equivalent to reducing the cortical layer thickness to 1/4 of the original one. In both the numerical and gel experimental results, it can be observed that small, tightly packed folds are formed. In Fig. 5**b**, we performed another modified numerical experiment with the cortical thickness doubled globally. We also performed a modified gel brain experiment by surface-coating 8 layers of PDMS gel onto the core layer instead of 4 layers as in the original gel model to double the cortical layer thickness. In both the numerical and gel experimental results, it can be observed that the number of small folds is significantly reduced. In both cases, the numerical and gel results show a good qualitative match.

**Figure 5.**
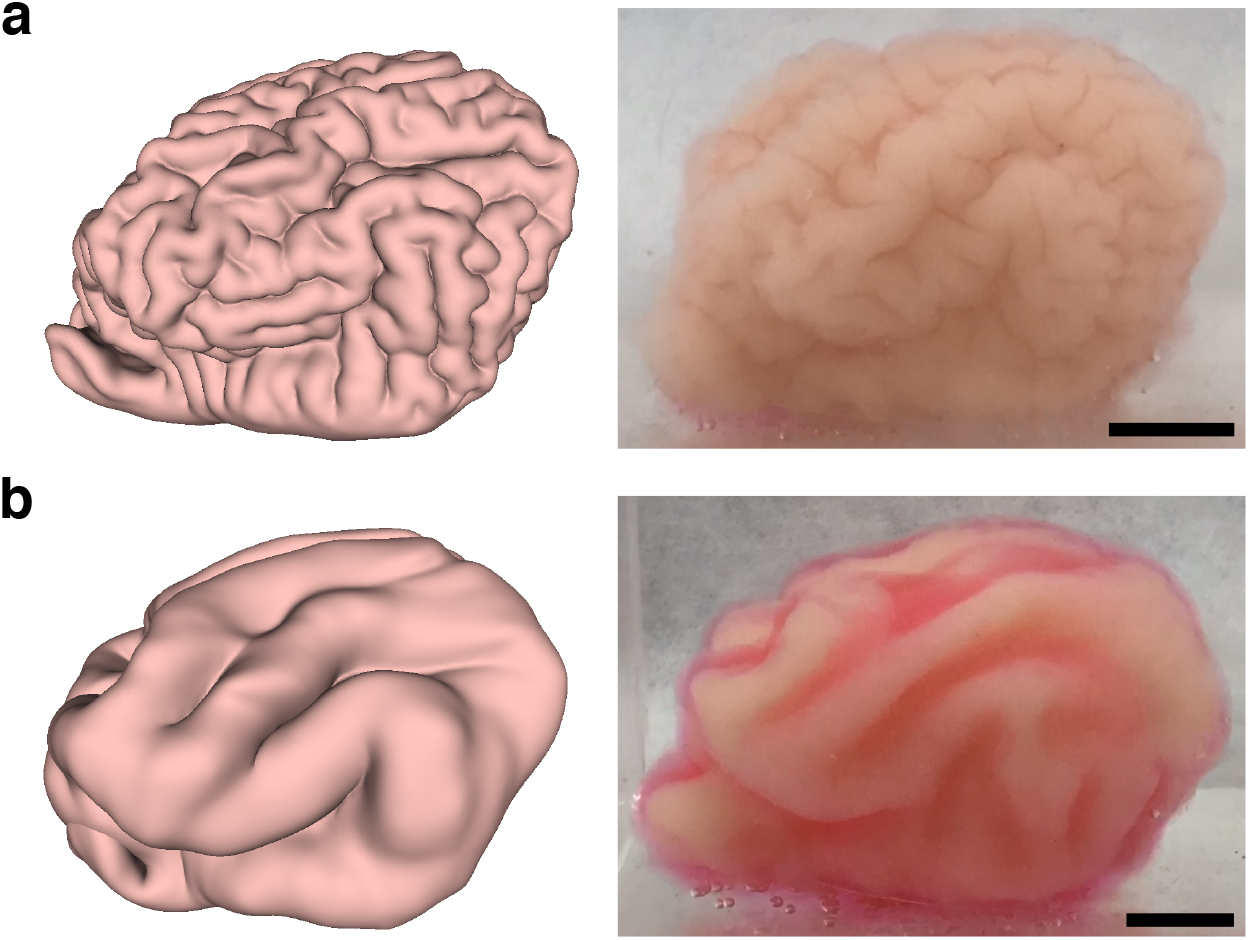
Numerical and gel experiments on the P8 ferret brain with globally modified brain gyrification. **a**, The numerical and gel experimental results with a global reduction of the cortical thickness to 1/4 of the original thickness. **b**, The numerical and gel experimental results with a global increase in the cortical thickness to twice the original thickness. Scale bar = 1cm.

### Neurology of ferret and human cortical malformations

After performing the modified physical gel and computational experiments for ferret brain misfoldings, we proceed to consider the effect of human genetic variants on brain gyrification in ferrets in prior studies (see Table 1). For instance, *SCN3A* encodes a sodium channel and specific missense mutations of it are associated with the MCD polymicrogyria (PMG) ***Smith et al. (2018)***, while genetic manipulations of *ASPM* have also been shown to produce severe microcephaly in human and ferret brains ***Johnson et al. (2018)***, while disrupting *TMEM161B* leads to cortical malformations in humans and ferrets ***Akula et al. (2023b)***. In these and other examples, various genetic causes lead to variations in the cortical thickness ratio *h*/*R* and/or the tangential growth ratio *g* in space and time, which we know to be critical geometric parameters that change the physical nature of the sulcification instability driving cortical folding.

**Table 1.** Human genetic variants modeled in ferret developmental brain phenotypes.

| Gene | Change in geometry | Relevant brain developmental disorder |
| --- | --- | --- |
| *SCN3A* *Smith et al. (2018)* | Excessive number of tightly packed folds | Polymicrogyria |
| *ASPM* *Johnson et al. (2018)* | Reduced brain size and cortical surface area | Microcephaly |
| *Cdk5* *Shinmyo et al. (2017)* | Reduced depth of the sulcus | Lissencephaly |
| *ARHGAP11B* *Kalebic et al. (2018)* | Expansion in both the radial and tangential dimensions | Megalencephaly |
| *TMEM161B* *Akula et al. (2023b)* | Reduced gyrus size and sulcal depth | Lissencephaly |

In Fig. 6**a**, we show a control human brain MRI (top), a wild-type P16 ferret brain MRI (middle), and the stepwise numerical simulation result under the normal parameter setup (bottom). In Fig. 6**b**, (top) we show the MCD polymicrogyria (PMG) phenotype in humans associated with over-expression of a mutated *SCN3A* gene ***Smith et al. (2018)***. In Fig. 6**b** (middle), we show that the same mutation in ferrets leads to an increased number of tightly packed folds in the perisylvian region ***Smith et al. (2018)***. To model this cortical malformation, we performed a modified numerical simulation with the cortical thickness reduced to 1/4 of the original thickness at a localized zone around the perisylvian region of the P8 brain model. From the modified numerical simulation result (Fig. 6**b**, bottom), we can see that our model can qualitatively capture perisylvian PMG (see also SI Section S4).

**Figure 6.**
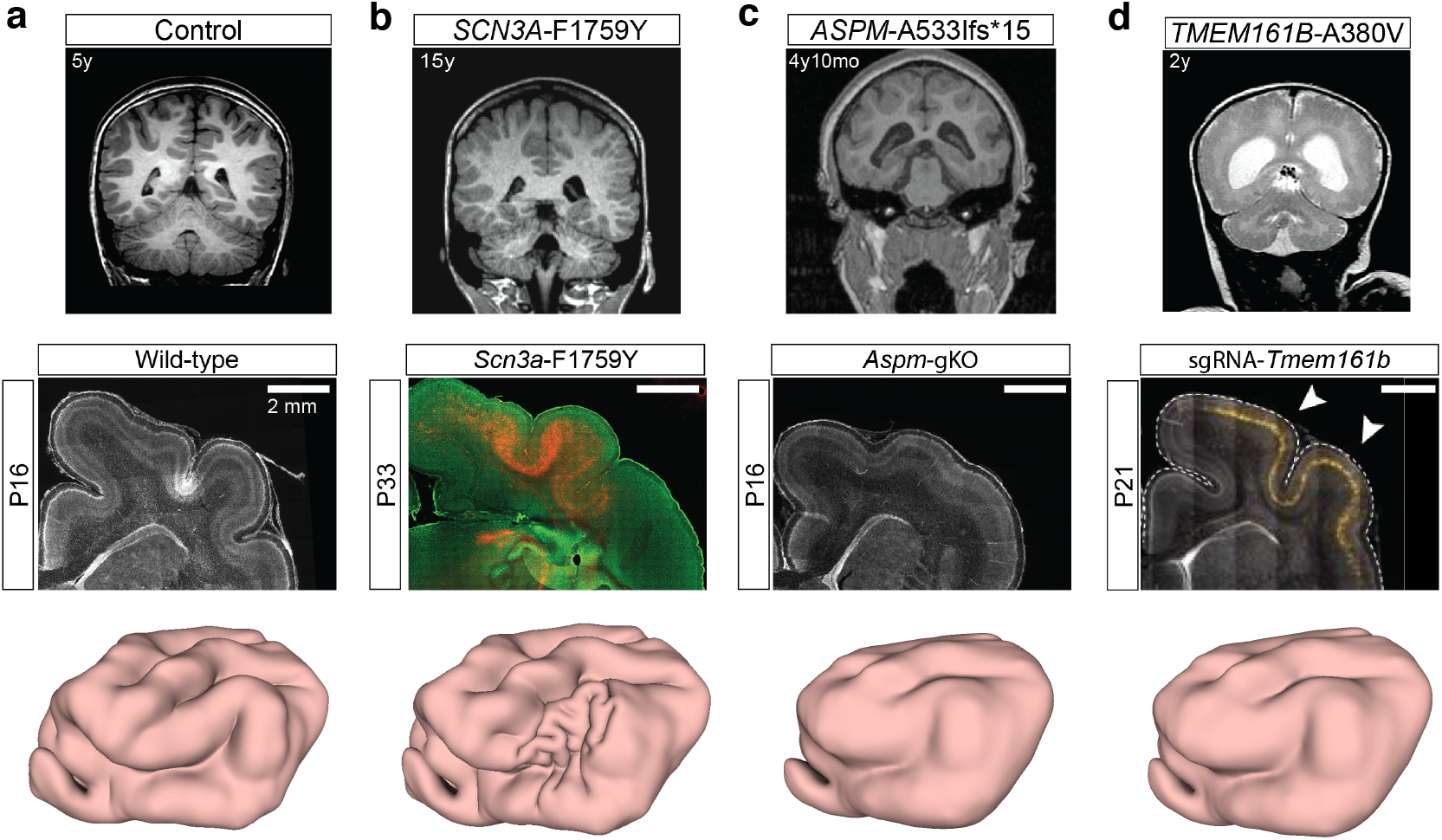
Modeling malformations of cortical development (MCD) using our model. **a**, Control. **b**, *SCN3A*. **c**, *ASPM*. **d**, *TMEM161B*. For each example, we show human (top) and ferret (middle) brain MRIs (images adapted from ***Smith et al. (2018); Johnson et al. (2018); Akula et al. (2023b)***). We then perform a modified numerical brain simulation on the P8 model with different tangential growth rate and cortical thickness parameters, including (**a**) the original growth rate and cortical thickness, (**b**) a reduction of the cortical thickness at a localized zone, (**c**) a reduction of the growth rate globally, and (**d**) a reduction of the growth rate and an increase of the cortical thickness globally. All numerical simulation results (bottom) qualitatively capture the cortical malformations.

In Fig. 6**c** (top) and Fig. 6**c** (middle) ***Johnson et al. (2018)*** we show that *ASPM* mutants produce severe microcephaly in human and ferret brains, because the cortical surface area is reduced while there is no significant change in the cortical thickness. To reproduce this malformation, we consider a modified computational experiment with a reduction of the growth rate to 1/4 of the original rate. In Fig. 6**c** (bottom), we show the results of numerical simulations that lead to less prominent folding when compared to the normal brain, consistent with observations in human and ferret brains.

Finally, in Fig. 6**d** (top) and Fig. 6**d** (middle) ***Akula et al. (2023b)***, we show that disrupting *TMEM161B* also leads to cortical malformations in human and ferret brains, with shallower sulci. Modifying the morphogenetic simulations with a reduction of the growth rate to 3/4 of the original rate and an increase of the cortical thickness to 1.5 times the original thickness leads to results shown in Fig. 6**d** (bottom) that match the experimentally observed malformation patterns qualitatively.

In SI Section S4, we present additional computational experiments to demonstrate the effect of different combinations of the growth rate and cortical thickness parameters on the cortical malformation results.

## Discussion

Understanding the growth and form of the cerebral cortex is a fundamental question in neurobiology, and the experimentally accessible progressive postnatal development of the ferret brain makes it an ideal system for analysis. Here, we have used a combination of physical and computational models based on differential growth to show how ferret brain morphologies arise. One may also use the physical and computational models to compare the ferret brain folding patterns with those in macaques and humans, as in our companion study ***Yin et al. (2025)***.

By modifying the scaled cortical layer thickness and the tangential growth profile in our model, we have qualitatively reproduced various cortical malformations and shown how developmental mechanisms lead to morphological manifestations with potential functional implications. As several diseases, such as ion channels ***Smith and Walsh (2020); Smith et al. (2021)***, converge on brain malformations, future studies to validate across disease pathways could leverage these results. All together, our study elucidates the normal and abnormal folding in the ferret brain as a function of its genetic antecedents that lead to changes in the geometry of the cortex and thence to different physical folding patterns with functional consequences. A computational and physical-gel brain study informed by detailed MRI of ferret and human fetal brains allows us to move towards a synthesis of the genetic, physical and morphological basis for cortical malformations. Natural next steps include accounting for varying spatio-temporal expansion rates of the cortex to capture the quantitative differences in the development of fetal folding patterns in both ferrets and humans, and understanding the functional consequences as a result of impaired connectivity due to misfolding.

## Methods

### Physical gel model for ferret brain folding

Beginning with T2-weighted motion-corrected anatomical MR images of ferret brains of various ages ***Toro et al. (2018)***, digital maps of the surfaces of native (pre-swollen) brain were recreated. Then, we followed our prior experimental approach ***Tallinen et al. (2014***, 2016) and produced two-layer PDMS gel models of the ferret brain at various ages based on the reconstructed brain surfaces. Specifically, we first generated a negative rubber mold with Ecoflex 00-30 from a 3D-printed brain plastic model and then the core gel with SYLGARD 184 at a 1:45 crosslinker:base ratio. To mimic the cortical layer, we surface-coated 4 layers of PDMS gel at a 1:35 crosslinker:base ratio onto the core layer. Finally, tangential cortical growth was mimicked by immersing the two-layer gel brain model in n-hexane for 1.5 hours, which resulted in solvent-driven swelling of the outer layers, leading to folding patterns. See SI Section S1 for details.

### Computational model for ferret brain folding

Three geometrical parameters of the 3D brain models control its morphogenesis: the average brain size *R* (determined for example by its volume), the average cortical thickness *T* and the average tangential expansion ratio of the cortex relative to the white matter, *g*^2^. To characterize brain development in the ferret, we followed the empirical scaling laws for gray-matter volume to thickness described in ***Tallinen et al. (2014)*** and set *R*/*T* ≈ 10 with the tangential expansion ratio *g* ≈ 1.9, along with an indicator function *θ*(*y*) = (1+*e*^10(*y*/*T* −1)^)^−1^, with *y* the distance from surface in a material reference frame used to distinguish between the cortical gray matter layer (with *θ* = 1) from the deeper white matter (with *θ* = 0).

Using MRIs of ferret brains, we created a computational model of the initial brain size and shape, which was discretized using tetrahedral meshes with over one million tetrahedral elements using Netgen ***NGSolve Team (2019)***. Using a finite element method implemented using a discretized version of the energy of the system (1), we minimized the energy by quasistatic equilibration using an explicit solver ***Tallinen et al. (2016)***, while growth was applied incrementally using the form described earlier by expanding the tetrahedral elements with inversion handling ***Stomakhin et al. (2012)*** and a nodal pressure formulation ***Bonet and Burton (1998)***. Self-avoidance of the surface was handled using the penalty-based vertex-triangle contact processing ***Ericson (2004)***. We also enforce the condition that there is no growth in the central part as well as in the bottom part of the brain to better simulate the development of ferret brains. See SI Section S2 for more details.

### Malformations of cortical development

In Fig. 6, we showed various human and ferret brain MRIs with malformations of cortical development. The *SCN3A* MRI images were adapted from ***Smith et al. (2018)*** (human: 15y; ferret: P33). The *ASPM* MRI images were adapted from ***Johnson et al. (2018)*** (human: 4y 10 mo; ferret: P16). The *TMEM161B* MRI images were adapted from ***Akula et al. (2023b)*** (human: 2y; ferret: P21).

The numerical simulation results for cortical malformations were obtained using the computational model described above, with the cortical thickness and growth rate parameters modified either locally or globally. See SI Section S4 for more details.

## Data Availability Statement

Requests for the ferret brain data should be made to ***Toro et al. (2018)***. All other data are included in the article and/or supplementary material.

## Code Availability Statement

Computer codes for numerical simulations and morphometric analyses are available on GitHub at https://github.com/garyptchoi/ferret-brain-morphogenesis.

## Acknowledgments

We thank Roberto Toro and Katja Heuer for the ferret brain data, and Jun Young Chung and James Weaver for their help with preliminary experiments. G.P.T.C. and L.M. are supported in part by the Harvard Quantitative Biology Initiative and the NSF-Simons Center for Mathematical and Statistical Analysis of Biology at Harvard, award no. 1764269. G.P.T.C. is also supported by the CUHK Faculty of Science Direct Grant for Research (Project Code 4053650). R.S.S. is supported by R00NS112604 and R01NS140046. C.A.W. is supported by the NINDS through R01NS032457 and R37NS035129, grant 62587 from the John Templeton Foundation (the opinions expressed in this publication are those of the authors and do not necessarily reflect the views of the John Templeton Foundation), and by the Allen Discovery Center for Human Brain Evolution, a Paul G. Allen Frontiers Group advised program of the Paul G. Allen Family Foundation. C.A.W. is an Investigator of the Howard Hughes Medical Institute. L.M. is also supported by the Simons Foundation and the Henri Seydoux Fund.

## Supplementary Material

## S1 Physical gel model for ferret brain folding

### S1.1 Surface reconstruction

Surfaces of initial (pre-swelling) brain states were reconstructed from T2-weighted motion-corrected anatomical MR images of ferret brains at various ages [1, 2] by R. Toro and K. Heuer. The reconstruction can be achieved by first computing masks of the pial and inner cortical surfaces from MRI data using the Nilearn Python module to generate and threshold image histograms [3], and then utilizing the Brainbox web application [4] to manually correct region of interest (ROI) selections and convert masks from Nifti (.nii.gz) files to three-dimensional triangular meshes, with conservative corrections to the output meshes including smoothing and inversion of inward-pointing normal vectors applied, with further mesh corrections performed using Meshlab.

### S1.2 Swelling experiment

In main text Fig. 2, we presented the swelling experimental results for an input P8 gel model and an input P16 gel model. In Fig. S1, we present the swelling experiment for the P4 ferret brain with the same experimental setup, where the 2-layer PDMS model was immersed in n-hexane for 1.5 hours. It can be observed that while the sulcal pattern is not particularly prominent in the input P4 gel model, complex folding patterns can be observed in the resulting swollen gel.

**Figure S1.**
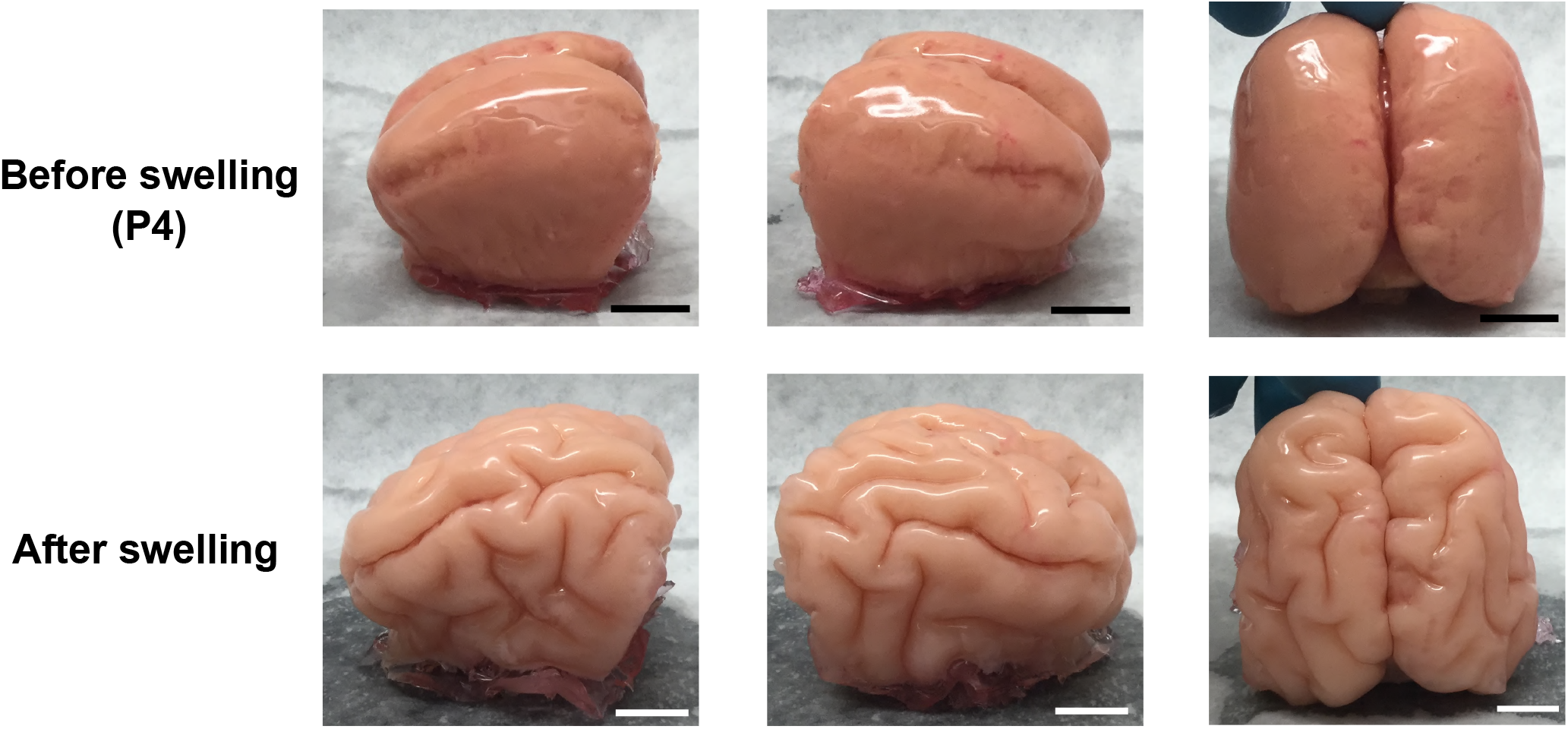
Swelling experiment for an input P4 gel brain. Here, the two-layer PDMS model of a P4 ferret brain was immersed in n-hexane for 1.5 hours. Analogous to the gel brain experiments presented in the main text, gyrification patterns can be observed in the swollen gel brain. Scale bar = 1cm.

We remark that comparing the P4 swelling result with the P8 and P16 gel brain experiments in main text Fig. 2, one can see that the folding patterns are notably different. A possible reason is that all gel models were immersed in n-hexane for swelling with the same duration (1.5 hours) regardless of their size and shape, which might result in the variation in the final gyrification patterns.

## S2 Computational model for ferret brain folding

### S2.1 Formulation

For the numerical simulation of ferret brain development, we follow the approach in [5, 6] and consider a material consisting of a layer of gray matter on top of a deep layer of white matter. The material is assumed to be neo-Hookean with volumetric strain energy density

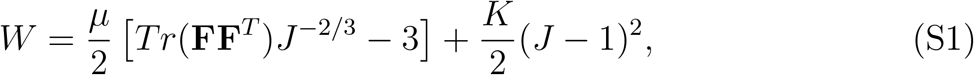

where **F** is the deformation gradient, *J* = det(**F**), *µ* is the shear modulus and *K* is the bulk modulus. We assume that *K* = 5*µ* for a modestly compressible material.

Three geometrical parameters of the 3D brain models are the brain size *R*, the cortical thickness *T* and the tangential expansion *g*^2^. For ferret brain development, we follow the empirical scaling law for gray-matter volume to thickness and set *R/T* ≈ 10 with the tangential expansion ratio *g* ≈ 1.9. An indicator function

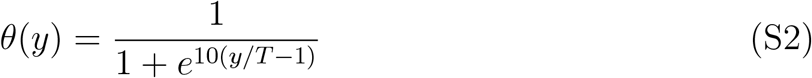

is applied for distinguishing between the cortical layer (with *θ* = 1) and white matter zone (with *θ* = 0). Here, *y* is the distance from surface in material coordinates.

### S2.2 Numerical experiments

Tetrahedral meshes were generated from the MRI-based reconstructed surfaces using the Netgen software [7], with each of them consisting of over 1 million tetrahedral elements. The numerical simulation procedure was implemented in C++. The energy of the system was minimized by quasistatic equilibration using an explicit scheme. Growth was applied by expanding the tetrahedral elements with inversion handling [8] and nodal pressure formulation [9]. We further set the innermost part and the bottom part of the brain tetrahedral meshes to be non-growing regions to better simulate the development of ferret brains. See [5, 6] for more details of the computation setup.

We simulated the stepwise growth of the ferret brain from P0 to P4, from P4 to P8, from P8 to P16, and from P16 to P32. Fig. S2 shows different views of the simulation results.

**Figure S2.**
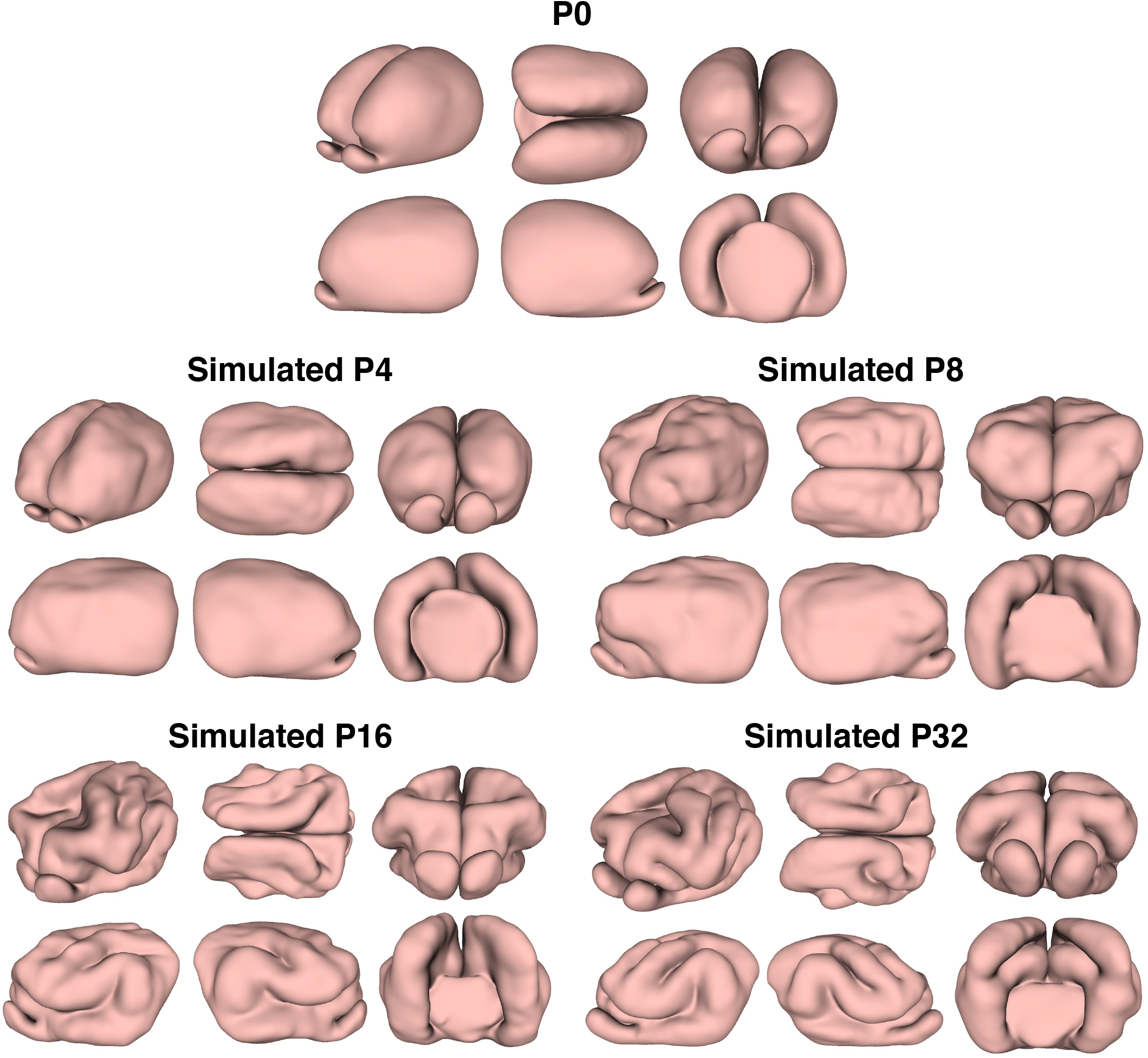
Stepwise growth simulation of ferret brain. The P0 brain and the simulated brains from P0 to P4, from P4 to P8, from P8 to P16, and from P16 to P32 are shown. For each brain, six different views are provided (not to scale).

**Figure S3.**
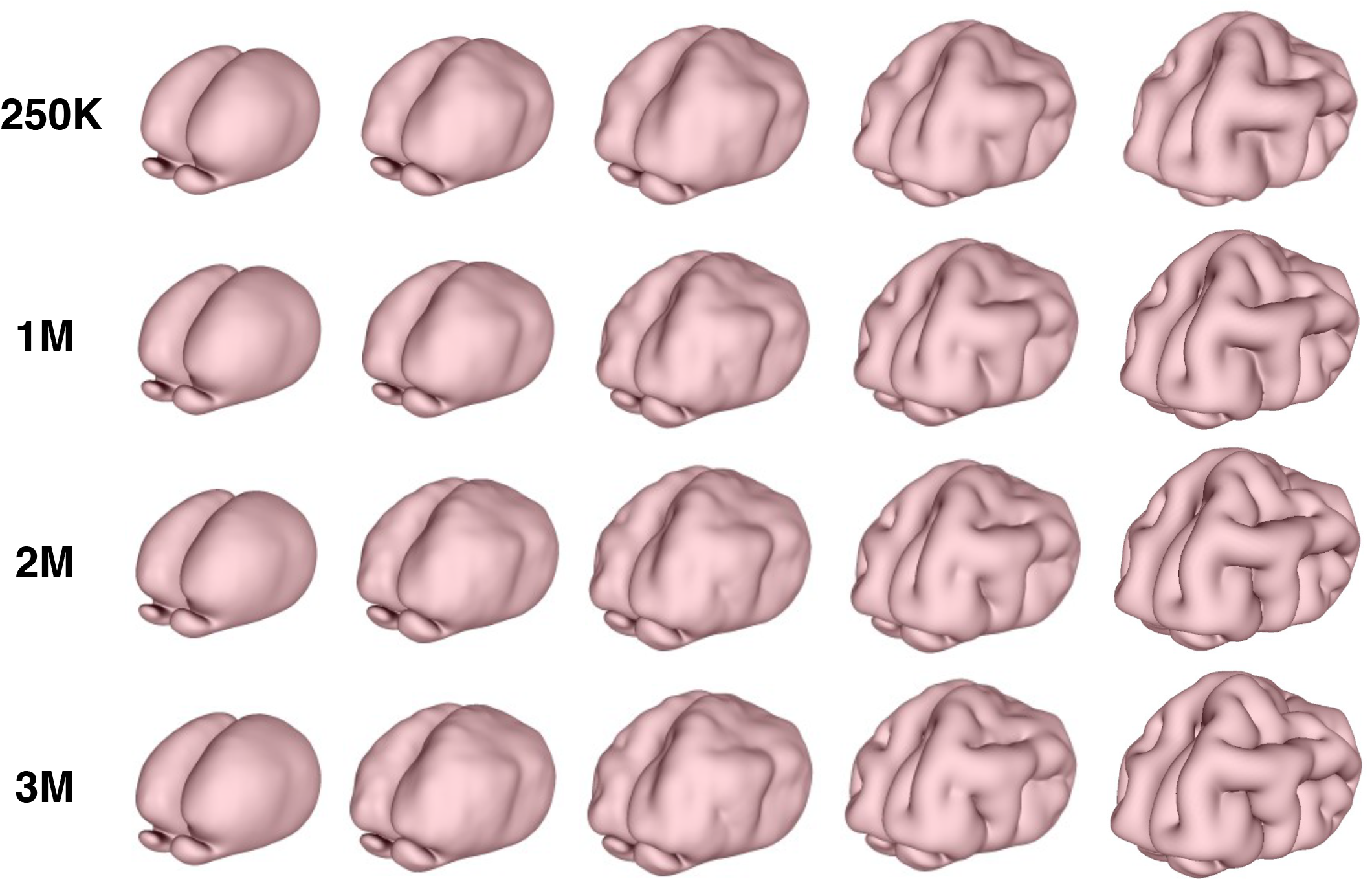
Mesh independence for the numerical simulation. The continuous growth simulations from P0 to adolescence using P0 brain tetrahedral meshes with 250,000 tets (first row), 1 million tets (second row), 2 million tets (third row), and 3 million tets (fourth row). It can be observed that the resulting folding patterns are highly similar.

### S2.3 Mesh independence

One may ask about the effect of mesh size on the numerical simulation. Fig. S3 shows the continuous simulation results starting from P0 with different mesh resolutions. It can be observed that the results are similar, which shows the mesh independence of the numerical simulation.

## S3 Morphometric analysis

### S3.1 Analyzing the brain surfaces using surface parameterization

The FLASH (Fast Landmark-Aligned Spherical Harmonic Parameterization) method [10] was used for quantifying the similarity of the numerically simulated brain and the MRI-based brain. Denote the two brain surfaces to be compared by *ℳ* and *N*. The FLASH method produces two bijective spherical mappings *f* : *ℳ* → *S*^2^ and *g* : *N* → *S*^2^, so that both surfaces are mapped to the unit sphere with prescribed landmark pairs on them optimally aligned. Here, *g* is a conformal map from *N* to the unit sphere *S*^2^, and *f* is a landmark-constrained optimized conformal map from *ℳ* to *S*^2^. More specifically, *f* achieves a balance between the landmark mismatch error and the conformality distortion, with the following combined energy minimized:

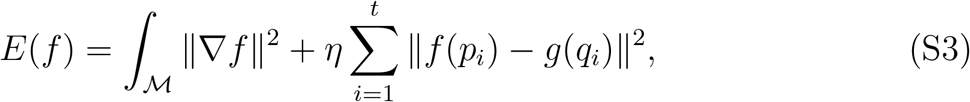

where {*p*_*i*_, *q*_*i*_} are corresponding landmark curves on *ℳ* and *N* respectively, and *η* is a balancing parameter. The coronal sulcus (cns), suprasylvian sulcus (sss), presylvian sulcus (prs), and pseudosylvian sulcus (pss) on both the left and right hemispheres of the ferret brain are used as the landmark curves to ensure that the two spherical parameterizations are consistent. We set *η* = 10 to achieve an accurate alignment of the two spherical parameterizations without inducing a large conformal distortion.

After obtaining the spherical parameterizations with the landmarks optimally aligned, we can compare the folding patterns of the two brain surfaces by evaluating the similarity of their shape indices on *S*^2^. The shape index is a dimensionless surface measure defined based on the surface mean curvature and the surface Gaussian curvature [11]. Specifically, the mean curvature *H* and Gaussian curvature *K* of any surface *S* are given by

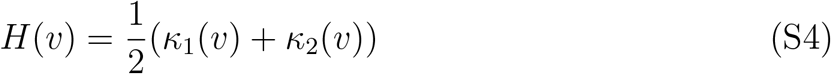

and

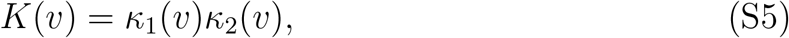

where *v* is a point on *S*, and *κ*_1_(*v*), *κ*_2_(*v*) are the principal curvatures at *v*. The shape index *I*(*v*) [11] at each point *v* is then given by

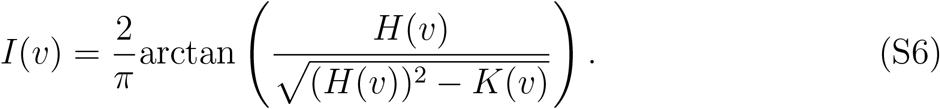

It is easy to see that *I*(*v*) ∈ [− 1, 1] for any *v*. Intuitively, for brain cortical surfaces, the sulci are with a smaller value of *I* and the gyri are with a larger value of *I*.

Now, with the aid of the spherical parameterizations, we can easily evaluate the similarity of the shape index distributions of two brain surfaces. Specifically, the similarity *s* of the shape index distributions *I*_*ℳ*_, *I*_*N*_ of the two surfaces *ℳ, N* is defined as

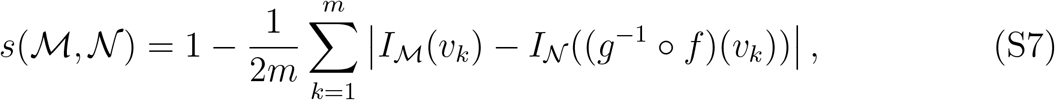

where 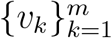 are the vertices of *ℳ*. In particular, since *I*_*ℳ*_, *I*_*N*_ ∈ [−1, 1], we have

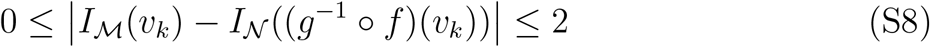

for all *k*. Consequently, we have *s* ∈ [0, 1], and *s* = 1 if the two brain surfaces are identical. As discussed in the main text, our result of *s* ≈ 0.83 shows that the real and simulated brain surfaces are highly similar.

Besides analyzing the entire brain surfaces, one can also quantify the shape difference for each half-brain surface using the mapping and shape similarity quantification method and different norms as described in [12]:

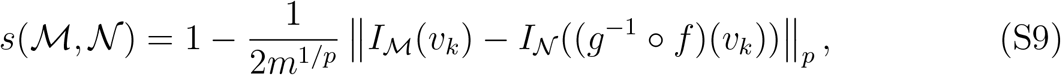

where ∥ · ∥_*p*_ is the *p*-norm. As shown by the quantification results in Table S1, the real and simulated half-brain surfaces are highly similar. These additional results again demonstrate the effectiveness of our model for simulating ferret brain folding.

### S3.2 Spherical harmonics representations

With the spherical parameterizations, we further computed the spherical harmonics representation of the two surfaces with different maximum order *N* used. Denote (*θ, ϕ*) as the spherical coordinates of *S*^2^, where *θ* ∈ [0, *π*] is the elevation angle and *ϕ* ∈ [−*π, π*] is the azimuth angle. The spherical harmonics functions 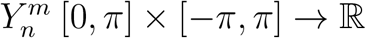of order *n* and degree *m* are given as follows [13]:

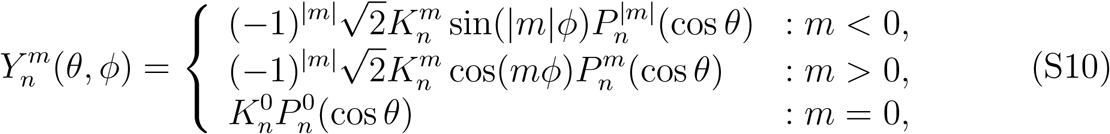

where

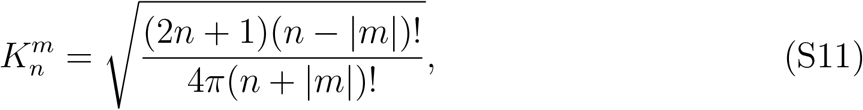

and

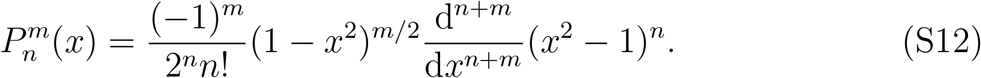

With the maximum order *N* prescribed, we can consider all 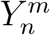 with *n* = 0, 1, …, *N* and *m* = − *n*, −*n* + 1, …, 0, …, *n* − 1, *n*. We can then approximate any given surface as a linear combination of this set of SH functions:

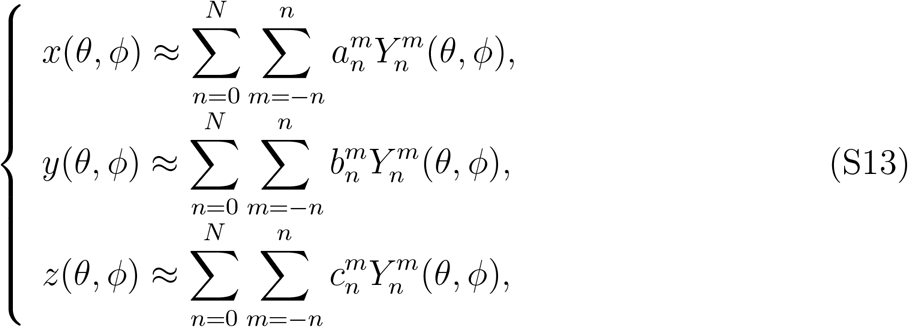

where the coefficients 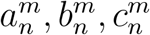 can be computed from the spherical parameterization (see [14] for details). As the maximum order *N* increases, more geometric details of the surface can be captured in the spherical harmonics representation.

From Fig. S4, it can be observed that for different maximum order *N*, the spherical harmonics representations of MRI-based brain and the numerically simulated brain match very well. More specifically, when *N* = 5 is used, both representations give an overall smooth brain shape without folding. When *N* = 15 is used, both of them start to exhibit some folding at consistent locations. When *N* = 25 is used, both of them show highly similar sulci and gyri patterns. This again indicates that the two brain surfaces are geometrically similar.

**Figure S4.**
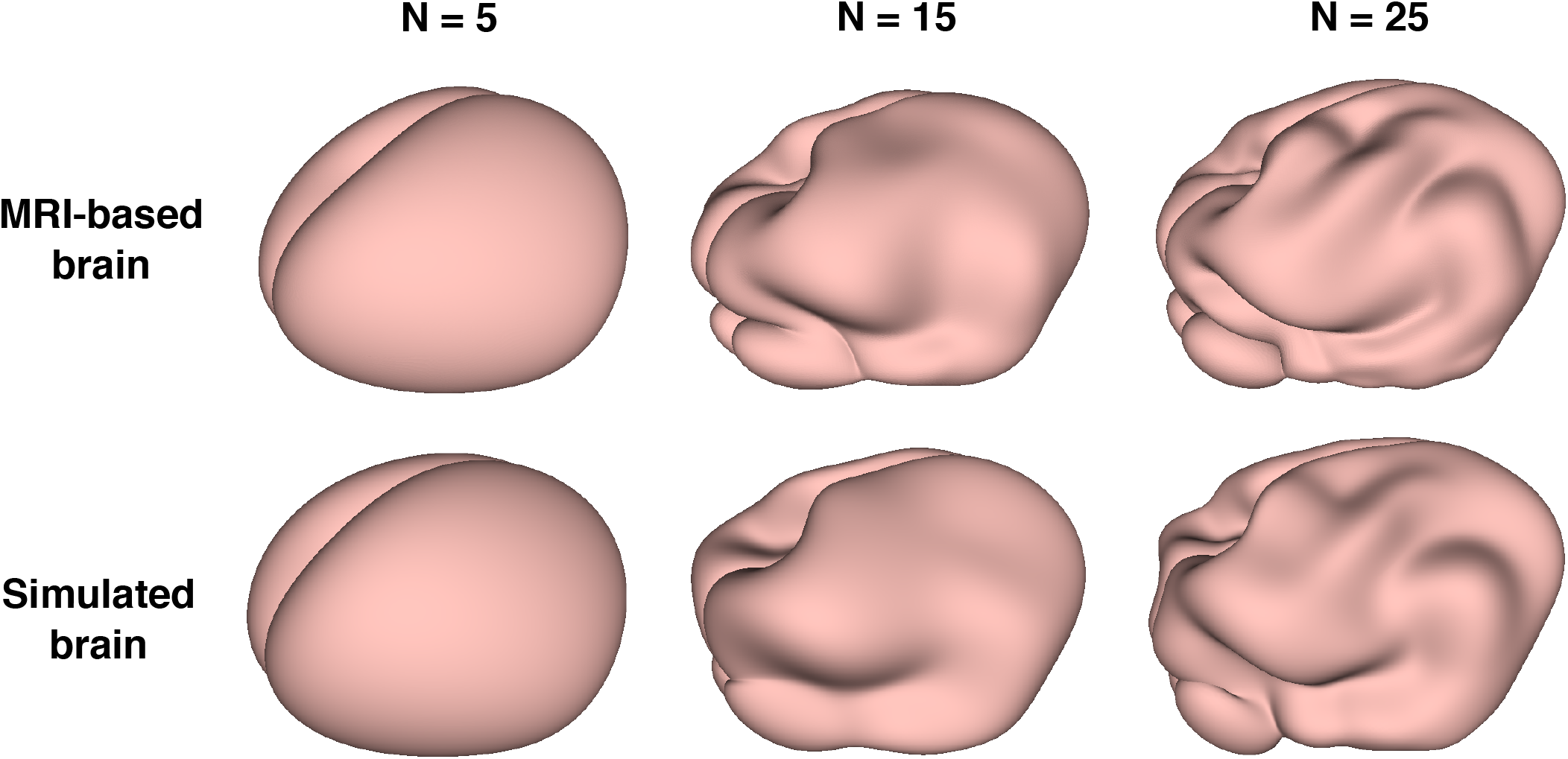
The spherical harmonics representations of the MRI-based brain and the numerically simulated brain with different maximum order *N* used.

## S4 Cortical malformations

In the main text, we considered various types of cortical malformations and discussed how our proposed model can be modified to capture their features qualitatively. In this section, additional analyses are provided.

In Fig. S5, we provide a comparison between human brain surface MRI reconstructions and our ferret brain simulations for the MCD polymicrogyria (PMG). Here, the human brain surface reconstruction images (*SCN3A* case and control age-matched subject) are adapted from [15], where MRI stacks were processed to extract the surfaces of the gray/white matter and gray matter/cerebrospinal fluid boundaries, and to define the cortical surface using FreeSurfer software. It can be observed that using our computational model with a modification in the cortical thickness, we obtain ferret brain simulation results with a good qualitative match.

**Figure S5.**
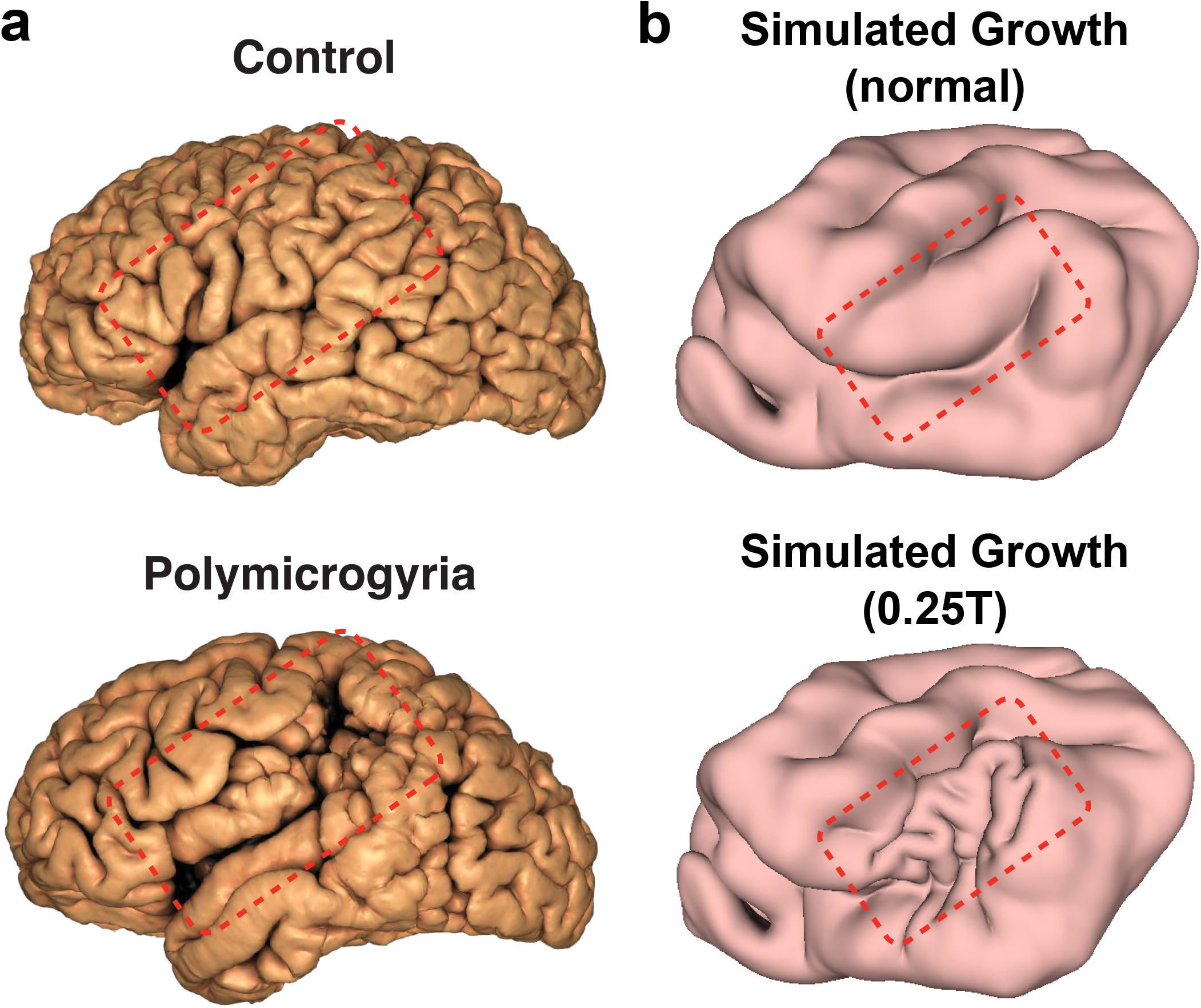
Comparison between human brain surface MRI reconstructions and our ferret brain simulations for the MCD polymicrogyria (PMG). **a**, Surface MRI reconstructions of human MRIs performed using FreeSurfer software of control (top) and age-matched affected individual with a gain-of-function *SCN3A* variant resulting in polymicrogyria. Red box outlines the PMG of perisylvian and surrounding areas, where the normal gyri and sulci form microgyri/sulci, pushing together to make an appearance of “Moroccan leather”. Images adapted from [15]. **b**, The numerical growth simulations on the P8 ferret brain without modification in the cortical thickness (top) and with a reduced cortical thickness to 1*/*4 of the original thickness, i.e., 0.25*T* (bottom). The differences are highlighted by the red boxes.

In Fig. S6, we further performed numerical simulations on the P8 ferret brain with different modifications in the tangential growth rate and cortical thickness at a localized region (dotted ellipse). Specifically, we first considered increasing the tangential growth rate at the localized region to 1.5 times or 2 times the original rate. In the numerical simulation results, more prominent folds can be observed at the corresponding region. Also, we considered reducing the cortical thickness to 1*/*2 or 1*/*4 of the original thickness at the localized region. In the simulation results, we can see that new folds are created at the localized region. Finally, we considered modifying both the tangential growth rate and the cortical thickness. In this case, a higher level of abnormal gyrification can be observed locally in the simulation results. Altogether, the modified brain experiments show that different cortical malformations can be effectively simulated by modifying the tangential growth rate and cortical thickness.

**Figure S6.**
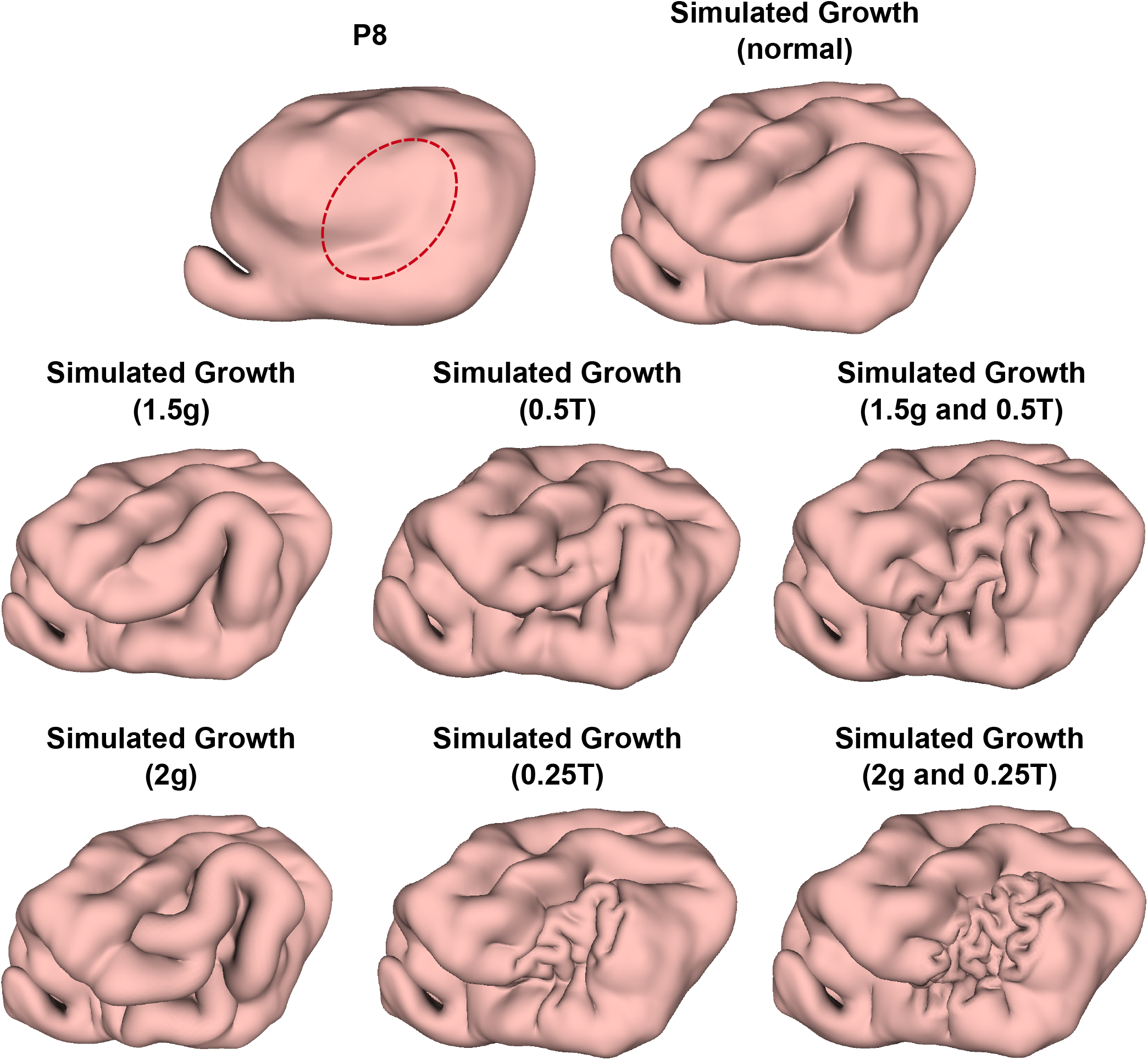
Numerical simulations with locally modified brain growth. We performed numerical simulations on the P8 brain with different modifications in the tangential growth rate *g* and the cortical layer thickness *T* at a localized region (dotted ellipse). The results without modification (normal), with an increased growth rate to 1.5 times the original growth rate (1.5*g*) or 2 times the original growth rate (2*g*), with a reduced cortical thickness to 1*/*2 of the original thickness (0.5*T*) or 1*/*4 of the original thickness (0.25*T*), and with a modification in both the growth rate and the cortical thickness (1.5*g* and 0.5*T*, 2*g* and 0.25*T*) are presented.

**Table S1.** Similarity between the real and simulated ferret half-brain surfaces.

| Surface | Similarity<br>(1-norm) | Similarity<br>(2-norm) | Similarity<br>(∞-norm) |
| --- | --- | --- | --- |
| Real P8 and<br>Simulated P8 (left) | 0.8040 | 0.7432 | 1.0000 |
| Real P8 and<br>Simulated P8 (right) | 0.7893 | 0.7260 | 1.0000 |
| Real P16 and<br>Simulated P16 (left) | 0.8247 | 0.7632 | 1.0000 |
| Real P16 and<br>Simulated P16 (right) | 0.7980 | 0.7339 | 1.0000 |

## S5 Video captions

**Video S1:** Time-lapse video of the swelling experiment for the P8 gel brain, where the 2-layer PDMS model was immersed in n-hexane for 1.5 hours.

**Video S2:** Time-lapse video of the swelling experiment for a P16 gel brain, where the 2-layer PDMS model was immersed in n-hexane for 1.5 hours.

**Video S3:** Continuous numerical simulation of the ferret brain folding from P0 to P32.

**Video S4:** Stepwise numerical simulations of the ferret brain folding from P0 to P4, from P4 to P8, from P8 to P16, and from P16 to P32.

## References

Akula SK, Exposito-Alonso D, Walsh CA. Shaping the brain: The emergence of cortical structure and folding. Dev Cell. 2023; 58(24):2836–2849.

Akula SK, Marciano JH, Lim Y, Exposito-Alonso D, Hylton NK, Hwang GH, Neil JE, Dominado N, Bunton-Stasyshyn RK, Song JHT, et al. TMEM161B regulates cerebral cortical gyration, Sonic Hedgehog signaling, and ciliary structure in the developing central nervous system. Proc Natl Acad Sci. 2023; 120(4):e2209964120.

Azevedo FAC, Carvalho LRB, Grinberg LT, Farfel JM, Ferretti REL, Leite REP, Filho WJ, Lent R, Herculano-Houzel S. Equal numbers of neuronal and nonneuronal cells make the human brain an isometrically scaled-up primate brain. J Comp Neurol. 2009; 513(5):532–541.

Bayly P, Okamoto R, Xu G, Shi Y, Taber L. A cortical folding model incorporating stress-dependent growth explains gyral wavelengths and stress patterns in the developing brain. Phys Biol. 2013; 10(1):016005.

Bonet J, Burton A. A simple average nodal pressure tetrahedral element for incompressible and nearly incompressible dynamic explicit applications. Comm Numer Meth Eng. 1998; 14(5):437–449.

Borrell V. How cells fold the cerebral cortex. J Neurosci. 2018; 38(4):776–783.

Budday S, Raybaud C, Kuhl E. A mechanical model predicts morphological abnormalities in the developing human brain. Sci Rep. 2014; 4:5644.

Choi PT, Lam KC, Lui LM. FLASH: Fast landmark aligned spherical harmonic parameterization for genus-0 closed brain surfaces. SIAM J Imaging Sci. 2015; 8(1):67–94.

Del-Valle-Anton L, Borrell V. Folding brains: from development to disease modeling. Physiol Rev. 2022; 102(2):511–550.

Desikan RS, Barkovich AJ. Malformations of cortical development. Ann Neurol. 2016; 80(6):797–810.

Ericson C. Real-time collision detection. CRC Press; 2004.

Fietz SA, Kelava I, Vogt J, Wilsch-Bräuninger M, Stenzel D, Fish JL, Corbeil D, Riehn A, Distler W, Nitsch R, et al. OSVZ progenitors of human and ferret neocortex are epithelial-like and expand by integrin signaling. Nat Neurosci. 2010; 13(6):690–699.

Garcia KE, Wang X, Kroenke CD. A model of tension-induced fiber growth predicts white matter organization during brain folding. Nat Commun. 2021; 12(1):6681.

Geschwind DH, Rakic P. Cortical evolution: judge the brain by its cover. Neuron. 2013; 80(3):633–647.

Gilardi C, Kalebic N. The ferret as a model system for neocortex development and evolution. Front Cell Dev Biol. 2021; 9:661759.

Hansen DV, Lui JH, Parker PRL, Kriegstein AR. Neurogenic radial glia in the outer subventricular zone of human neocortex. Nature. 2010; 464(7288):554–561.

Herculano-Houzel S. The human brain in numbers: a linearly scaled-up primate brain. Front Hum Neurosci. 2009; 3:857.

Heuer K, Traut N, de Sousa AA, Valk SL, Clavel J, Toro R. Diversity and evolution of cerebellar folding in mammals. Elife. 2023; 12:e85907.

His W. Untersuchungen über die erste Anlage des Wirbelthierleibes: die erste Entwickelung des Hühnchens im Ei, vol. 1. FCW Vogel; 1868.

Hohlfeld E, Mahadevan L. Unfolding the sulcus. Phys Rev Lett. 2011; 106(10):105702.

Hutton C, Draganski B, Ashburner J, Weiskopf N. A comparison between voxel-based cortical thickness and voxel-based morphometry in normal aging. Neuroimage. 2009; 48(2):371–380.

Johnson MB, Sun X, Kodani A, Borges-Monroy R, Girskis KM, Ryu SC, Wang PP, Patel K, Gonzalez DM, Woo YM, et al. Aspm knockout ferret reveals an evolutionary mechanism governing cerebral cortical size. Nature. 2018; 556(7701):370–375.

Kalebic N, Gilardi C, Albert M, Namba T, Long KR, Kostic M, Langen B, Huttner WB. Human-specific ARHGAP11B induces hallmarks of neocortical expansion in developing ferret neocortex. Elife. 2018; 7:e41241.

Koenderink JJ, van Doorn AJ. Surface shape and curvature scales. Image Vis Comput. 1992; 10(8):557–564.

Kroenke CD, Bayly PV. How forces fold the cerebral cortex. J Neurosci. 2018; 38(4):767–775.

Lui JH, Hansen DV, Kriegstein AR. Development and evolution of the human neocortex. Cell. 2011; 146(1):18–36.

Masuda K, Toda T, Shinmyo Y, Ebisu H, Hoshiba Y, Wakimoto M, Ichikawa Y, Kawasaki H. Pathophysiological analyses of cortical malformation using gyrencephalic mammals. Sci Rep. 2015; 5(1):1–15.

Molnár Z, Clowry GJ, Šestan N, Alzu’bi A, Bakken T, Hevner RF, Hüppi PS, Kostović I, Rakic P, Anton E, et al. New insights into the development of the human cerebral cortex. J Anat. 2019; 235(3):432–451.

Neal J, Takahashi M, Silva M, Tiao G, Walsh CA, Sheen VL. Insights into the gyrification of developing ferret brain by magnetic resonance imaging. J Anat. 2007; 210(1):66–77.

NGSolve Team, NetGen; 2019. https://ngsolve.org/.

Nie J, Guo L, Li G, Faraco C, Miller LS, Liu T. A computational model of cerebral cortex folding. J Theor Biol. 2010; 264(2):467–478.

Pang JC, Aquino KM, Oldehinkel M, Robinson PA, Fulcher BD, Breakspear M, Fornito A. Geometric constraints on human brain function. Nature. 2023; 618(7965):566–574.

Richman DP, Stewart RM, Hutchinson JW, Caviness VS. Mechanical model of brain convolutional development. Science. 1975; 189(4196):18–21.

Sawada K, Watanabe M. Development of cerebral sulci and gyri in ferrets (Mustela putorius). Congenit Anom. 2012; 52(3):168–175.

Schwartz E, Nenning KH, Heuer K, Jeffery N, Bertrand OC, Toro R, Kasprian G, Prayer D, Langs G. Evolution of cortical geometry and its link to function, behaviour and ecology. Nat Commun. 2023; 14(1):2252.

Shinmyo Y, Terashita Y, Duong TAD, Horiike T, Kawasumi M, Hosomichi K, Tajima A, Kawasaki H. Folding of the cerebral cortex requires Cdk5 in upper-layer neurons in gyrencephalic mammals. Cell Rep. 2017; 20(9):2131–2143.

Smith RS, Florio M, Akula SK, Neil JE, Wang Y, Hill RS, Goldman M, Mullally CD, Reed N, Bello-Espinosa L, et al. Early role for a Na+, K+-ATPase (ATP1A3) in brain development. Proc Natl Acad Sci. 2021; 118(25):e2023333118.

Smith RS, Kenny CJ, Ganesh V, Jang A, Borges-Monroy R, Partlow JN, Hill RS, Shin T, Chen AY, Doan RN, et al. Sodium channel SCN3A (NaV1. 3) regulation of human cerebral cortical folding and oral motor development. Neuron. 2018; 99(5):905–913.

Smith RS, Walsh CA. Ion channel functions in early brain development. Trends Neurosci. 2020; 43(2):103–114.

Stomakhin A, Howes R, Schroeder C, Teran JM. Energetically consistent invertible elasticity. In: Proceedings of the 11th ACM SIGGRAPH/Eurographics conference on Computer Animation; 2012. p. 25–32.

Tabata H, Nakajima K. Efficient in utero gene transfer system to the developing mouse brain using electroporation: visualization of neuronal migration in the developing cortex. Neuroscience. 2001; 103(4):865–872.

Tallinen T, Biggins JS. Mechanics of invagination and folding: Hybridized instabilities when one soft tissue grows on another. Phys Rev E. 2015; 92(2):022720.

Tallinen T, Biggins JS, Mahadevan L. Surface sulci in squeezed soft solids. Phys Rev Lett. 2013; 110(2):024302.

Tallinen T, Chung JY, Biggins JS, Mahadevan L. Gyrification from constrained cortical expansion. Proc Natl Acad Sci. 2014; 111(35):12667–12672.

Tallinen T, Chung JY, Rousseau F, Girard N, Lefèvre J, Mahadevan L. On the growth and form of cortical convolutions. Nat Phys. 2016; 12(6):588–593.

Toro R, Burnod Y. A morphogenetic model for the development of cortical convolutions. Cereb Cortex. 2005; 15(12):1900–1913.

Toro R, Perron M, Pike B, Richer L, Veillette S, Pausova Z, Paus T. Brain size and folding of the human cerebral cortex. Cereb Cortex. 2008; 18(10):2352–2357.

Toro R, Tiesinga P, Delzescaux T, Evans A, et al., FIIND: Ferret Interactive Integrated Neuro Development Atlas; 2018. https://neuroanatomy.github.io/fiind/.

Van Essen DC; Elsevier. Biomechanical models and mechanisms of cellular morphogenesis and cerebral cortical expansion and folding. Semin Cell Dev Biol. 2022; 140:90–104.

Welker W. Why does the cortex fissure and fold: a review of determinants of gyri and sulci. In: Cerebral Cortex (Jones EG, Peters A, eds) Plenum Press New York, NY; 1990. p. 3–136.

Yin S, Liu C, Choi GPT, Jung Y, Toro R, Heuer K, Mahadevan L. Morphogenesis and morphometry of brain folding patterns across species. eLife. 2025; 14:RP107138.

## References

[1] R. Toro, R. Bakker, T. Delzescaux, A. Evans, and P. Tiesinga, “FIIND: Ferret interactive integrated neurodevelopment atlas,” Res. Ideas Outcomes, vol. 4, p. e25312, 2018.

[2] “FIIND: Ferret interactive integrated neuro development atlas.” https://neuroanatomy.github.io/fiind/, 2018.

[3] A. Abraham, F. Pedregosa, M. Eickenberg, P. Gervais, A. Mueller, J. Kossaifi, A. Gramfort, B. Thirion, and G. Varoquaux, “Machine learning for neuroimaging with scikit-learn,” Front. Neuroinform., vol. 8, p. 14, 2014.

[4] R. Toro, “BrainBox github repository.” https://github.com/r03ert0/BrainBox, 2017.

[5] T. Tallinen, J. Y. Chung, J. S. Biggins, and L. Mahadevan, “Gyrification from constrained cortical expansion,” Proc. Natl. Acad. Sci., vol. 111, no. 35, pp. 12667–12672, 2014.

[6] T. Tallinen, J. Y. Chung, F. Rousseau, N. Girard, J. Lefévre, and L. Mahadevan, “On the growth and form of cortical convolutions,” Nat. Phys., vol. 12, no. 6, pp. 588–593, 2016.

[7] “NetGen.” https://ngsolve.org/, 2019.

[8] A. Stomakhin, R. Howes, C. Schroeder, and J. M. Teran, “Energetically consistent invertible elasticity,” in Proceedings of the 11th ACM SIGGRAPH/Eurographics Conference on Computer Animation, pp. 25–32, 2012.

[9] J. Bonet and A. Burton, “A simple average nodal pressure tetrahedral element for incompressible and nearly incompressible dynamic explicit applications,” Comm. Numer. Meth. Eng., vol. 14, no. 5, pp. 437–449, 1998.

[10] P. T. Choi, K. C. Lam, and L. M. Lui, “FLASH: Fast landmark aligned spherical harmonic parameterization for genus-0 closed brain surfaces,” SIAM J. Imaging Sci., vol. 8, no. 1, pp. 67–94, 2015.

[11] J. J. Koenderink and A. J. van Doorn, “Surface shape and curvature scales,” Image Vis. Comput., vol. 10, no. 8, pp. 557–564, 1992.

[12] S. Yin, C. Liu, G. P. T. Choi, Y. Jung, R. Toro, K. Heuer, and L. Mahadevan, “Morphogenesis and morphometry of brain folding patterns across species,” eLife, vol. 14, p. RP107138, 2025.

[13] C. Müller, Spherical harmonics, vol. 17. Springer, 2006.

[14] C. Brechbühler, G. Gerig, and O. Kübler, “Parametrization of closed surfaces for 3-D shape description,” Comput. Vis. Image Underst., vol. 61, no. 2, pp. 154–170, 1995.

[15] R. S. Smith, C. J. Kenny, V. Ganesh, A. Jang, R. Borges-Monroy, J. N. Partlow, R. S. Hill, T. Shin, A. Y. Chen, R. N. Doan, et al., “Sodium channel SCN3A (NaV1. 3) regulation of human cerebral cortical folding and oral motor development,” Neuron, vol. 99, no. 5, pp. 905–913, 2018.

